## Supplementary information for "Structural basis of substrate recognition and translocation by human ABCA4"

### Extended Data Figures

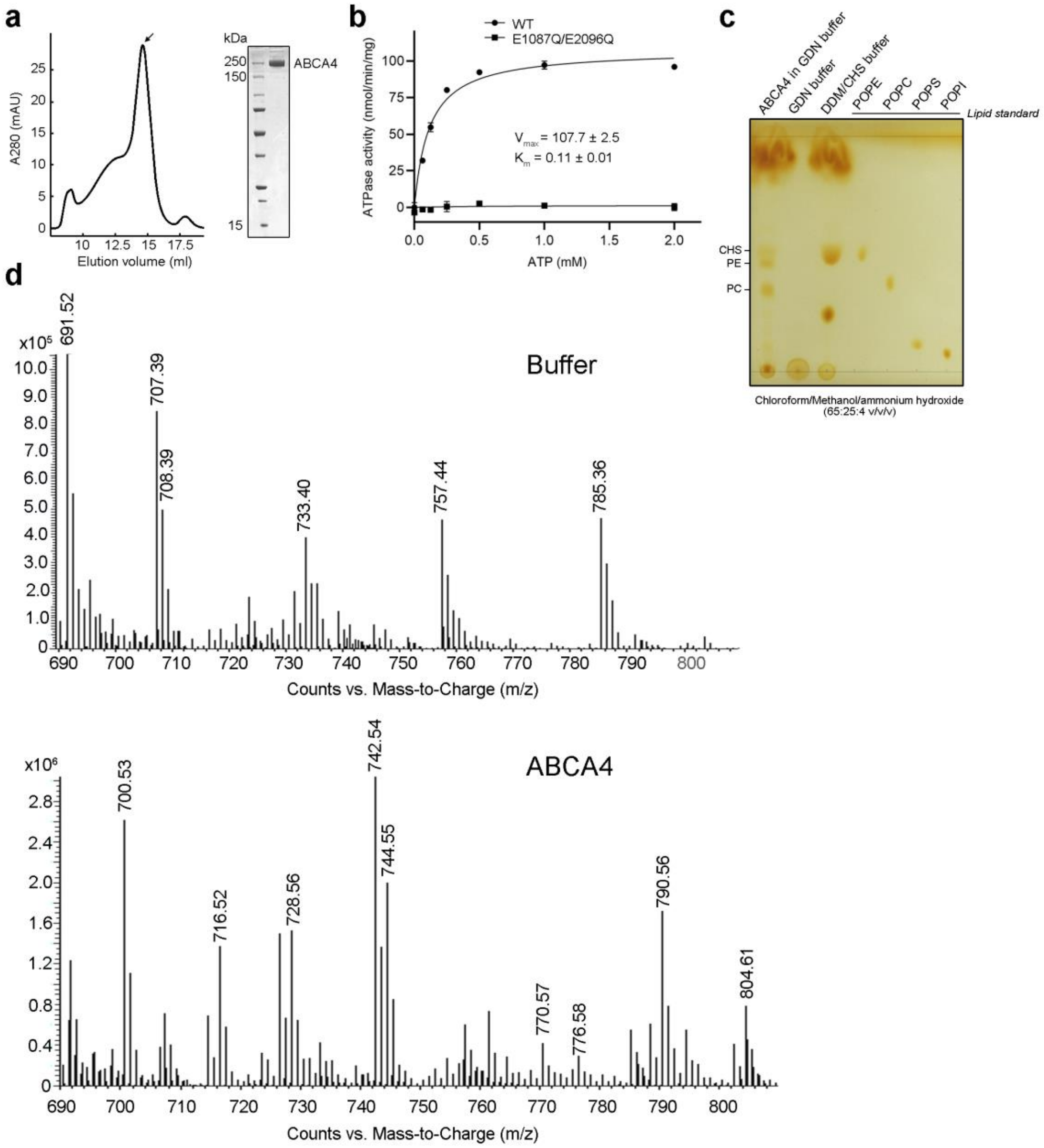


**Extended Data Fig. 1 | Biochemical characterizations of human ABCA4.** **a,** Size exclusion chromatography of human ABCA4. The peak fractions indicated by the arrow were pooled and concentrated for biochemical and structural studies. The purified protein was visualized by Coomassie-blue stained SDS-PAGE. **b,** Specific ATPase activity versus ATP concentration measured using wild-type (WT) ABCA4 protein or the catalytic mutant (E1087Q/E2096Q, $ABCA4_{QQ}$). **c,** Detection of endogenous lipids co-purified with ABCA4 by TLC. **d,** Mass spectrometry analysis of lipids co-purified with ABCA4. The upper panel represents the mass spectra of lipids purified from buffer (as a blank). The lower panel shows the mass spectra of lipids co-purified with ABCA4. The ions at *m/z* 770.53, 716.52, 728.56, 742.54, 744.55, and 770.57 were identified as PE. The ions at *m/z* 776.58, 790.56, and 804.61 were identified as PC. All the ions were confirmed by tandem MS spectra.

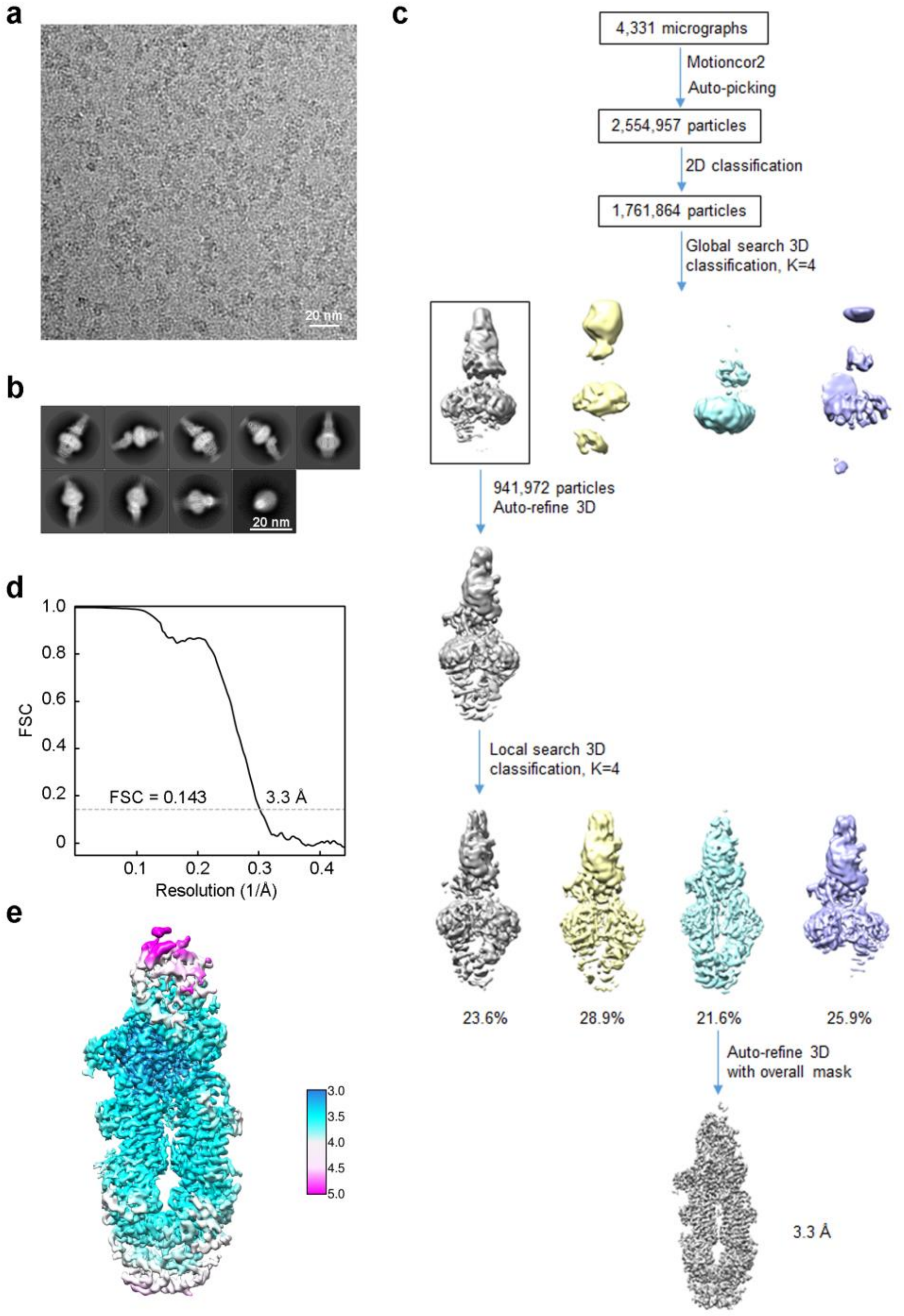


**Extended Data Fig. 2 | Cryo-EM analysis of apo ABCA4.** **a,** Representative cryo-EM micrograph. **b**, Representative 2D class averages. **c,** Flowchart for cryo-EM data processing. **d,** Gold-standard FSC curve for ABCA4 map generated using Relion 3.0. **e,** Local resolution map of ABCA4. The color code for resolutions, shown with the unit Å, is calculated using Relion 3.0.

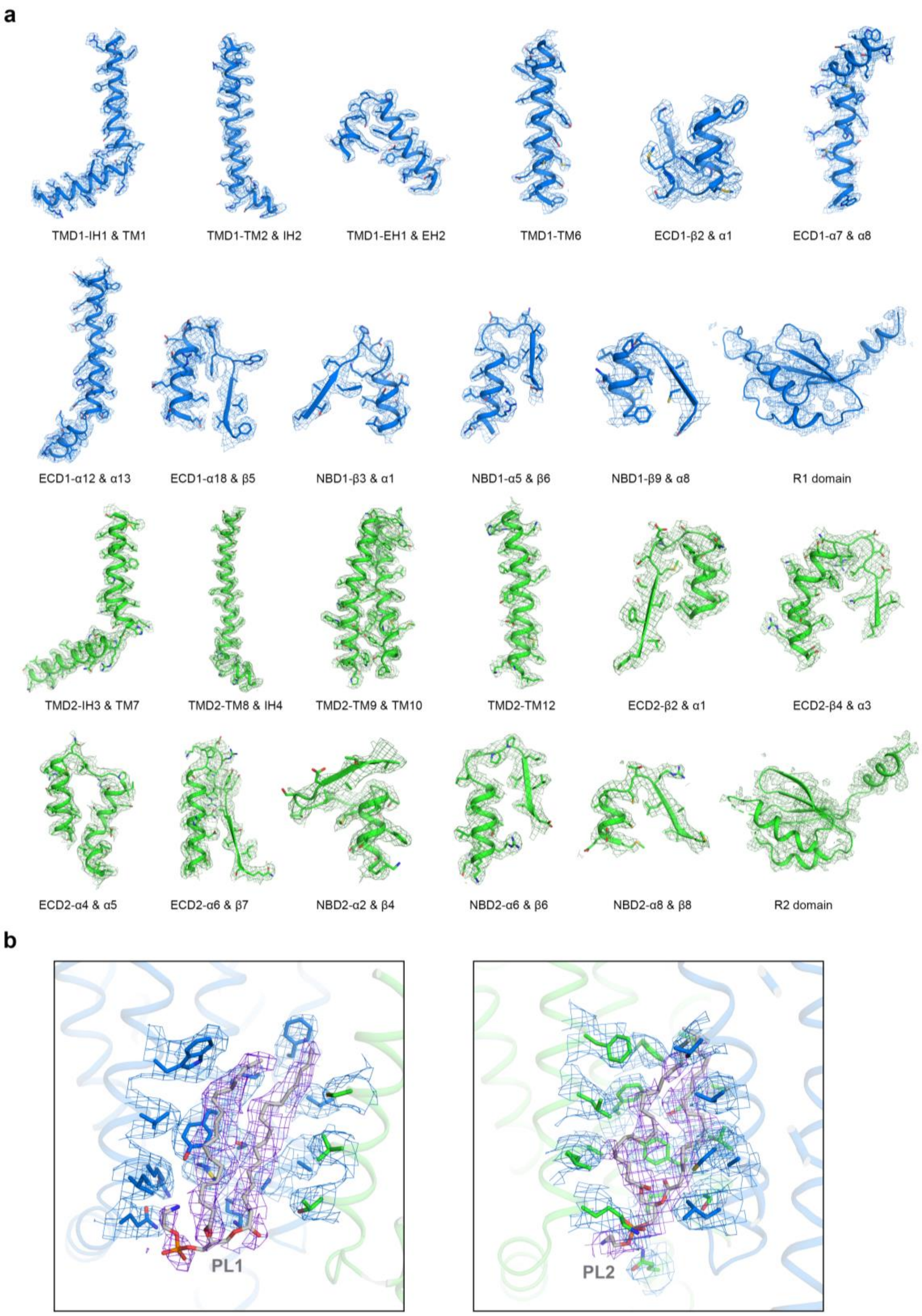


**Extended Data Fig. 3 | EM maps for representative segments (a) and phospholipid binding sites (b) of apo ABCA4.** Electron density maps, contoured at 5 σ (**a**) or 4 σ (**b**), were prepared in PyMol.

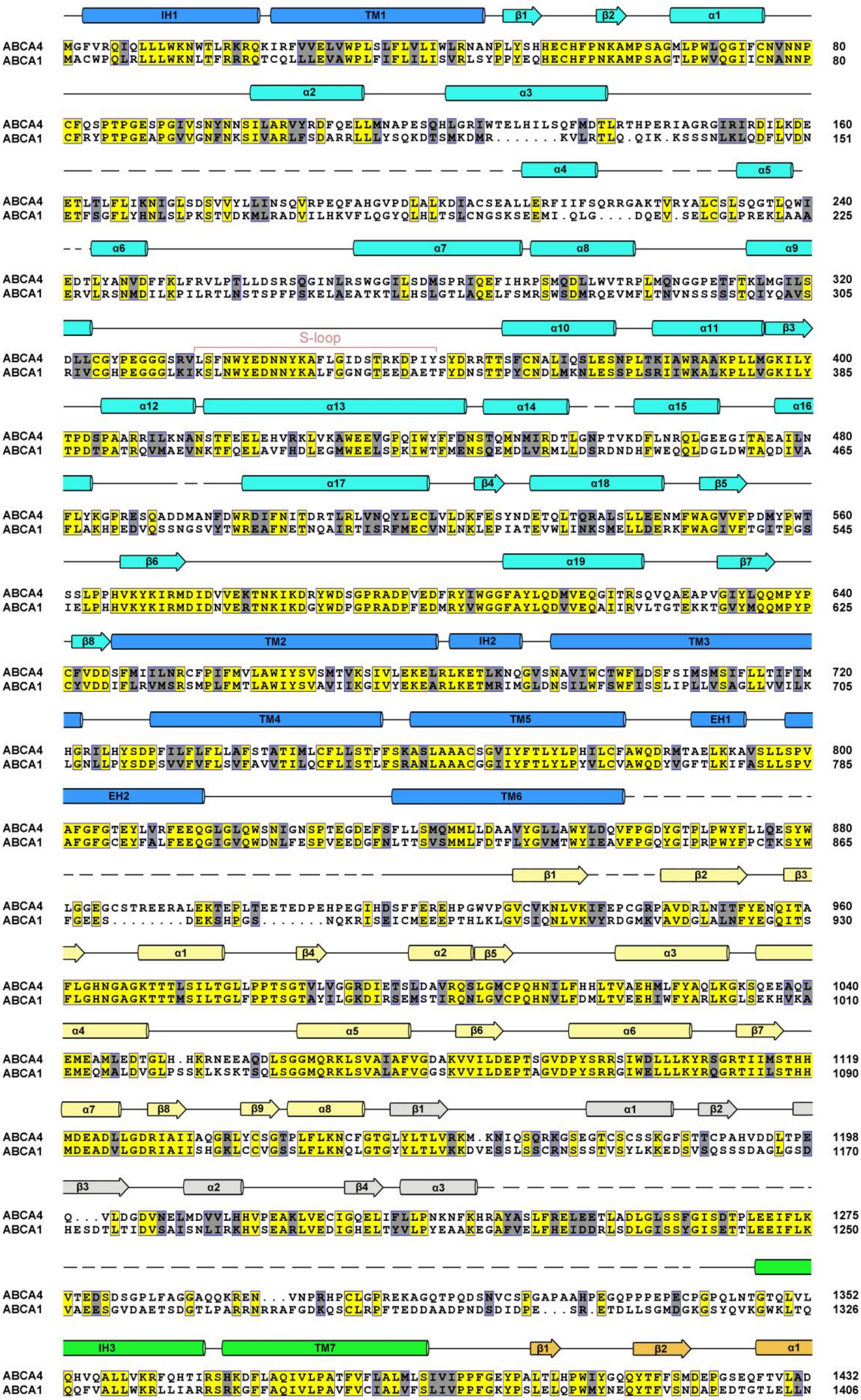
IH1
TM1
β1
β2
α1
ABCA4 80
ABCA1 80
α2
α3
ABCA4 160
ABCA1 151
α4
α5
ABCA4 240
ABCA1 225
α6
α7
α8
α9
ABCA4 320
ABCA1 305
S-loop
α10
α11
β3
ABCA4 400
ABCA1 385
α12
α13
α14
α15
α16
ABCA4 480
ABCA1 465
α17
β4
α18
β5
ABCA4 560
ABCA1 545
β6
α19
β7
ABCA4 640
ABCA1 625
β8
TM2
IH2
TM3
ABCA4 720
ABCA1 705
TM4
TM5
EH1
ABCA4 800
ABCA1 785
EH2
TM6
ABCA4 880
ABCA1 865
β1
β2
β3
ABCA4 960
ABCA1 930
α1
β4
α2
β5
α3
ABCA4 1040
ABCA1 1010
α4
α5
β6
α6
β7
ABCA4 1119
ABCA1 1090
α7
β8
β9
α8
β1
α1
β2
ABCA4 1198
ABCA1 1170
β3
α2
β4
α3
ABCA4 1275
ABCA1 1250
ABCA4 1352
ABCA1 1326
IH3
TM7
β1
β2
α1
ABCA4 1432
ABCA1 1406

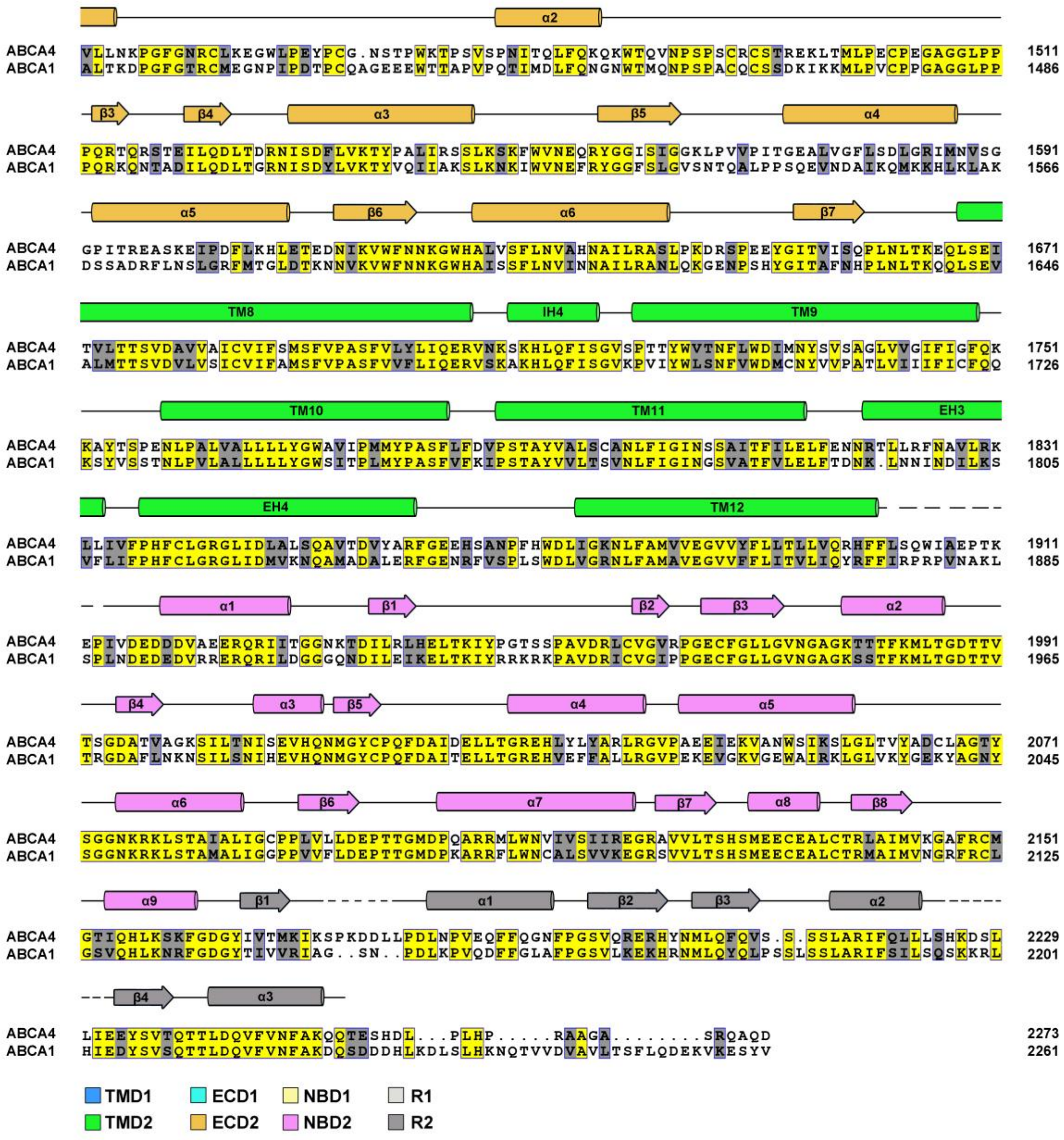

**Extended Data Fig. 4 | Sequence alignment of human ABCA4 and ABCA1.** Secondary structural elements of human ABCA4 are indicated above the sequence alignment. Dashed lines indicate protein portions that were not resolved in the cryo-EM map. Invariant and highly conserved amino acids are shaded yellow and gray, respectively. The secondary structural elements are colored based on the domains, with TMD1 and TMD2 colored marine and green; ECD1 and ECD2 colored cyan and orange; NBD1 and NBD2 colored yellow and magenta; R1 and R2 colored light gray and dark gray.

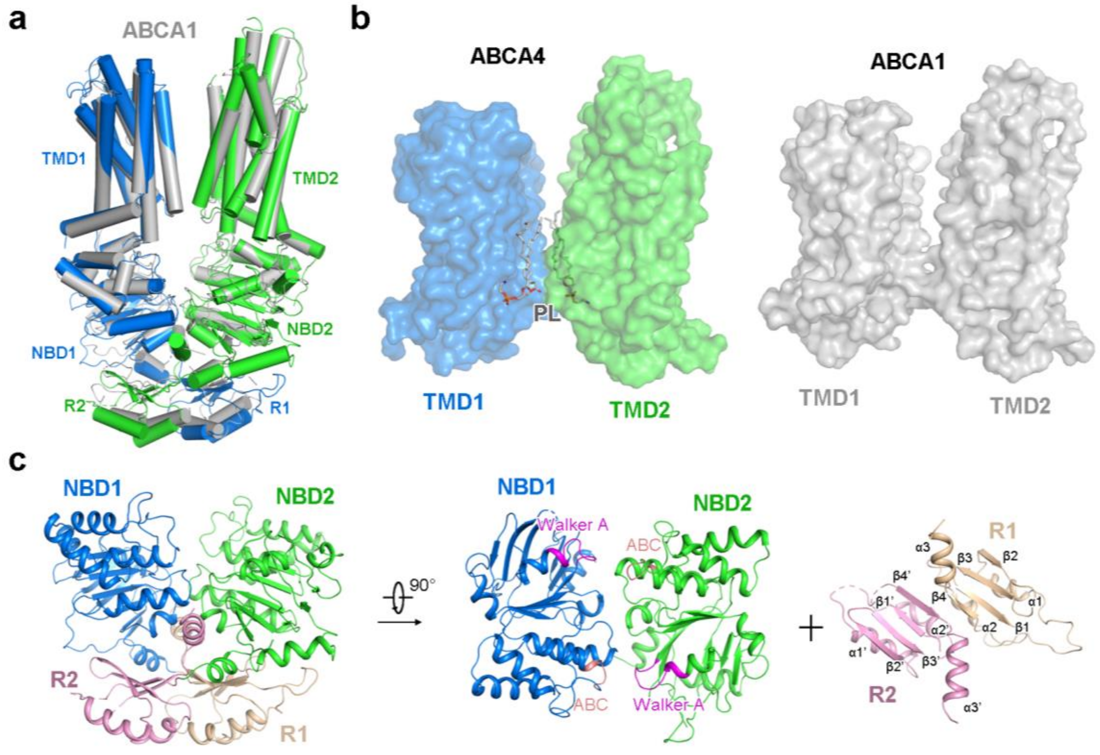


**Extended Data Fig. 5 | Structural features of the TMDs, NBDs, and RDs of ABCA4.** **a,** Superposition of ABCA4 and ABCA1 structures with ECDs omitted. ABCA1 structure is colored gray. TMD1, NBD1, and R1 of ABCA4 structure are colored marine. TMD2, NBD2, and R2 of ABCA4 structure are colored green. **b**, Surface representations of the TMDs from ABCA4 and ABCA1 structures. The phospholipids bound in the cytoplasmic leaflet of ABCA4 are shown as gray sticks. **c,** Structure of the NBDs and RDs of ABCA4. NBD1 and NBD2 are colored marine and green, respectively. R1 and R2 are colored wheat and pink, respectively. The Walker A motifs and ABC signature sequences are highlighted in magenta and light pink, respectively.

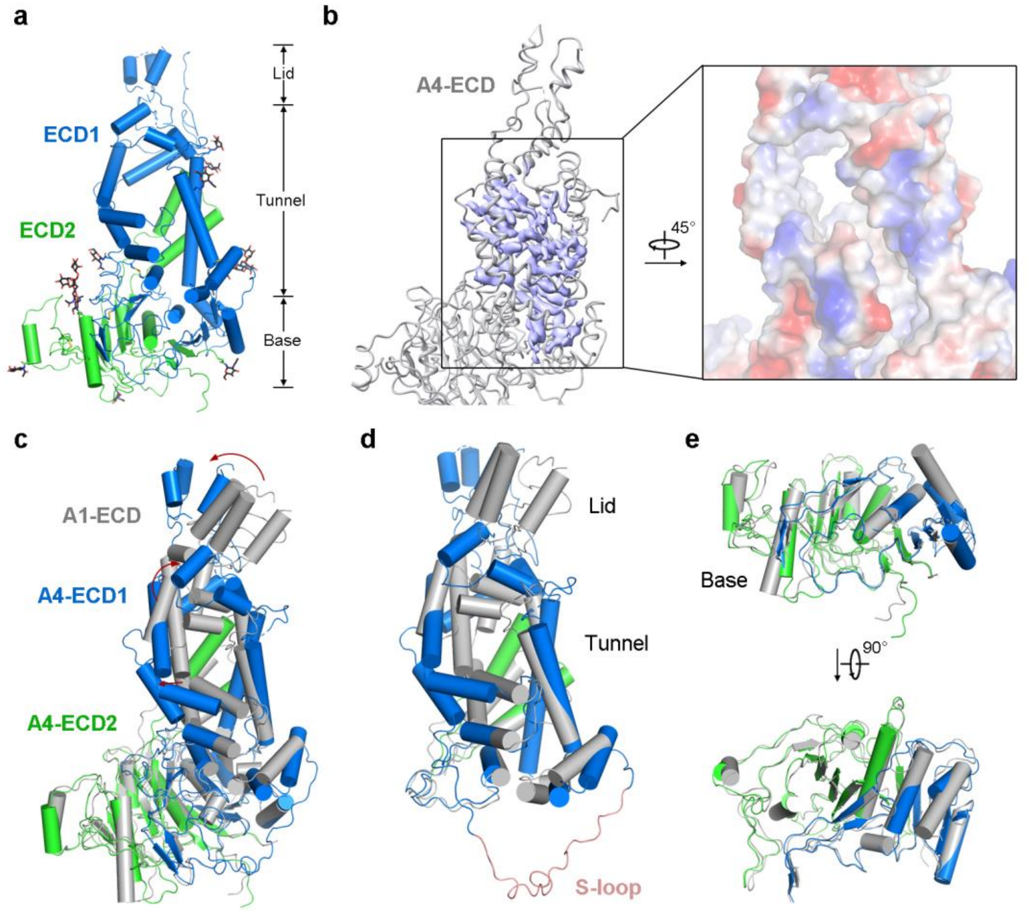


**Extended Data Fig. 6 | Structural features of the ECD of ABCA4. a,** The ECD of ABCA4 exhibits a three-layer structural organization, consisting of lid, tunnel, and base. **b**, A hydrophobic tunnel within the ECD of ABCA4 is filled with lipid-like densities. The lipid-like densities in the left panel are displayed in purple. The ECD tunnel in the right panel is shown as surface electrostatic potential. **c,** Superposition of the ECD structures from ABCA4 and ABCA1. The ECD from ABCA1 (A1-ECD) is colored gray. ECD1 and ECD2 of ABCA4 are colored marine and green, respectively. **d,** Structural differences in the tunnel and lid regions from A4-ECD and A1-ECD. The S-loop from A4-ECD tunnel is colored light pink. The corresponding region from A1-ECD tunnel was missing due to structural flexibility. **e,** Close-up views of the superimposed ECD bases.

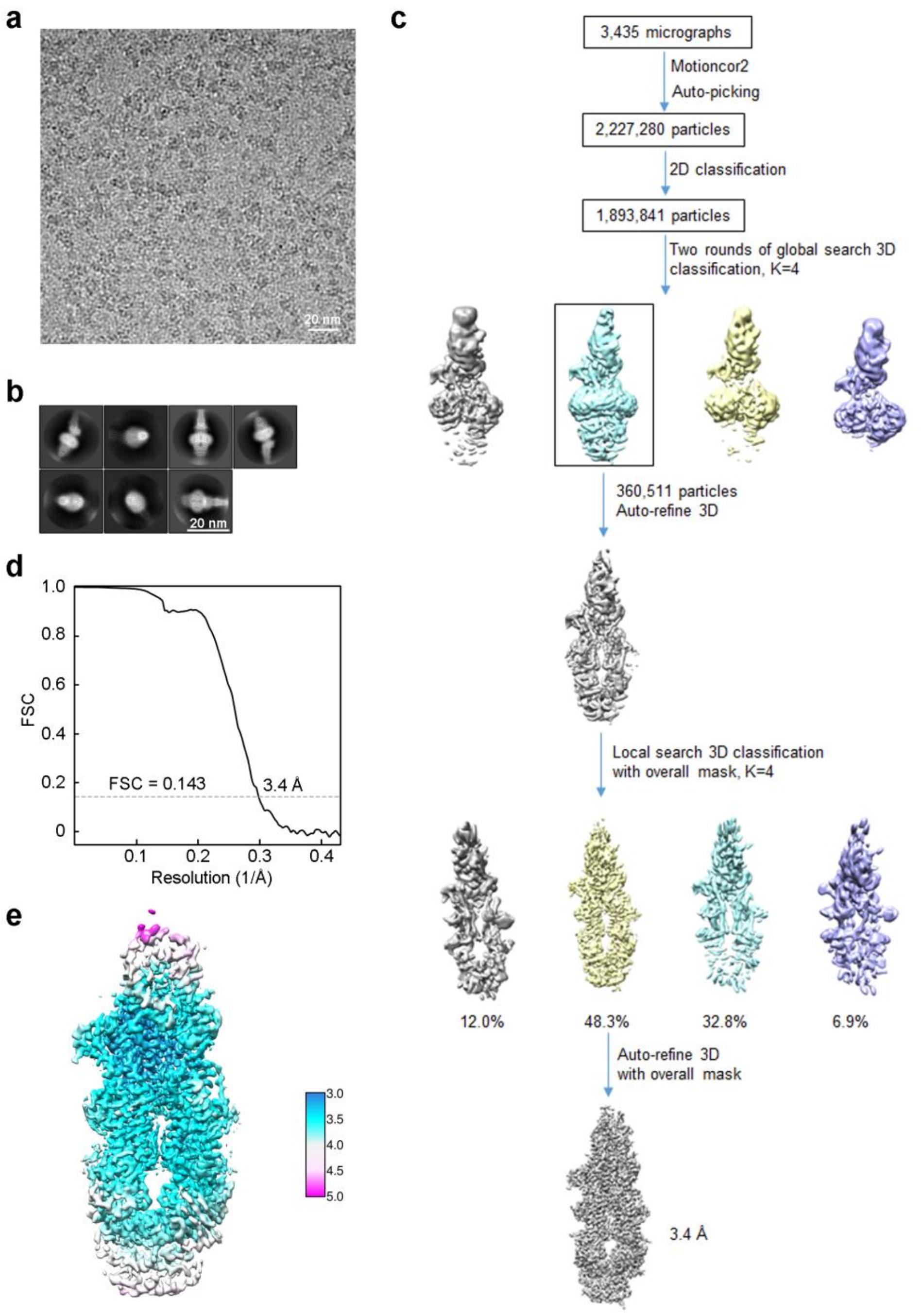


**Extended Data Fig. 7 | Cryo-EM analysis of NRPE-bound ABCA4.** **a,** Representative cryo-EM micrograph. **b**, Representative 2D class averages. **c,** Flowchart for cryo-EM data processing. **d,** Gold-standard FSC curve for NRPE-bound ABCA4 map. **e,** Local resolution map of NRPE-bound ABCA4.

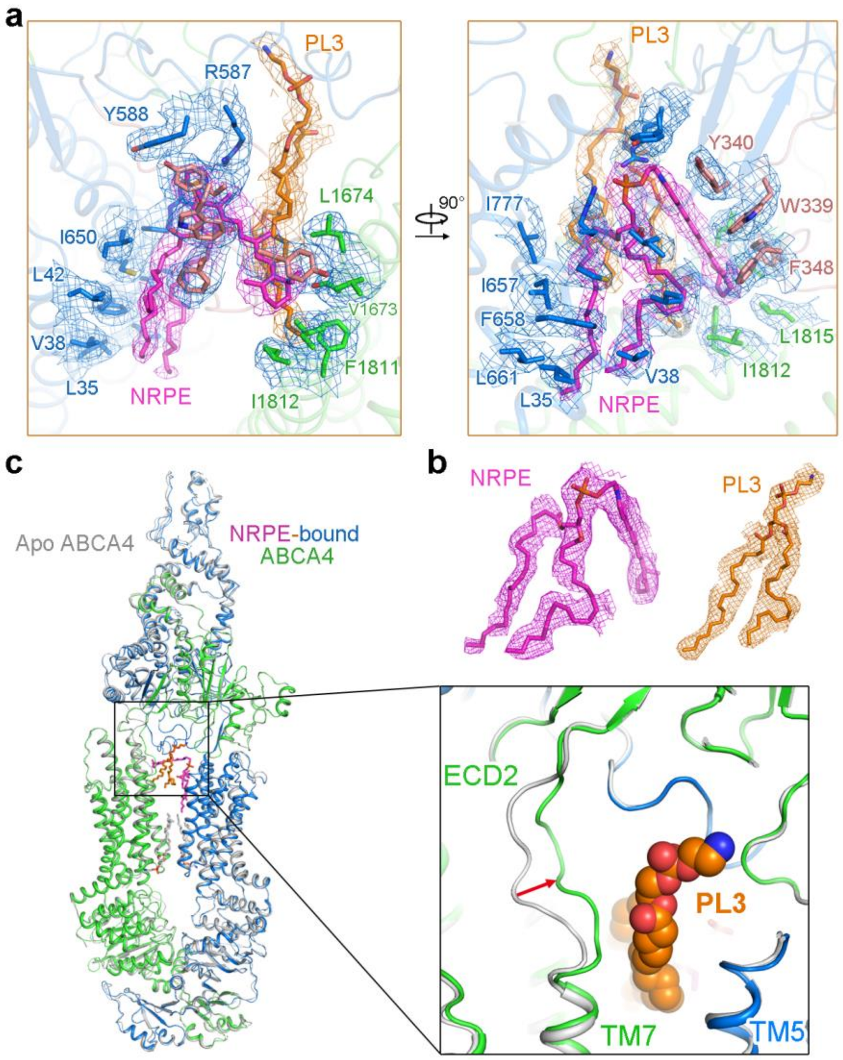


**Extended Data Fig. 8 | EM maps for NRPE binding site and a small conformational change of ABCA4 upon NRPE binding.** **a,** EM maps for the NRPE binding site contoured at 4 σ. **b**, EM maps for NRPE and PL3 bound in the luminal membrane leaflet contoured at 4 σ. **c,** A small conformational change of the connecting loop between TM7 and ECD2 upon NRPE and PL3 binding. Apo ABCA4 is colored gray. NRPE-bound ABCA4 is colored marine and green. PL3 is shown as orange spheres.

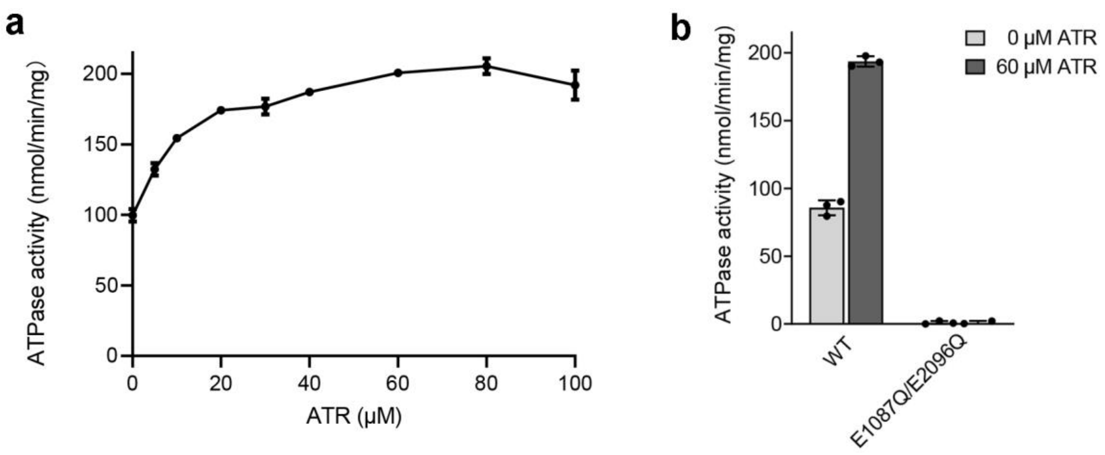


**Extended Data Fig. 9 | ATR stimulates the ATPase activity of ABCA4 purified in CHAPS/lipid mixture.** **a**, Specific ATPase activity of WT ABCA4 as a function of ATR concentration. **b**, Specific ATPase activity of WT ABCA4 or the catalytic mutant (E1087Q/E2096Q) in the absence or presence of 60 μM ATR.

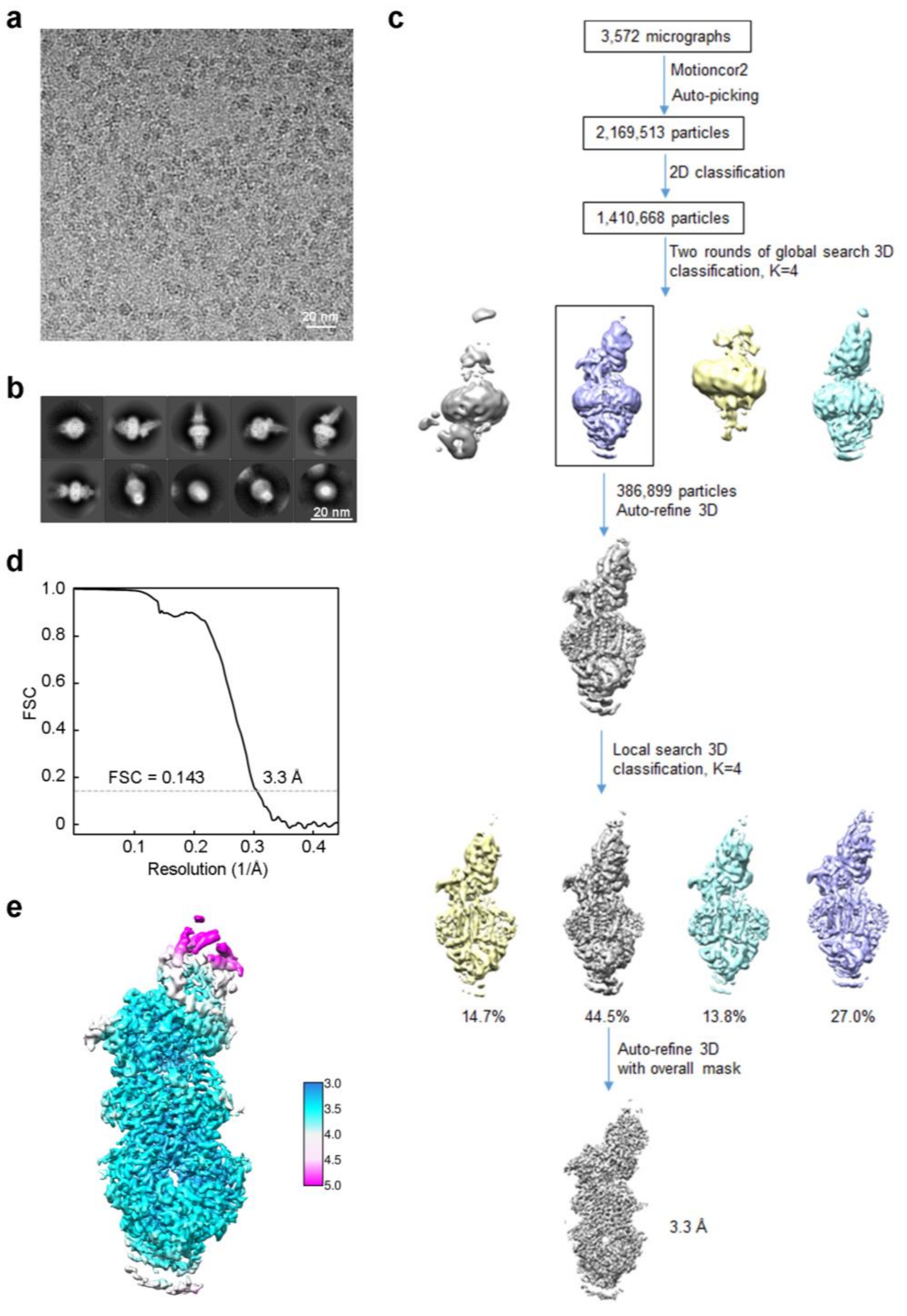


**Extended Data Fig. 10 | Cryo-EM analysis of ATP-bound ABCA4. a,** Representative cryo-EM micrograph. **b**, Representative 2D class averages. **c,** Flowchart for cryo-EM data processing. **d,** Gold-standard FSC curve for ATP-bound ABCA4 map. **e,** Local resolution map of ATP-bound ABCA4.

**Extended Data Table 1 | Cryo-EM data collection, refinement and validation statistics**

| | Apo ABCA4 (EMD-XXXX, PDB XXX) | NRPE-bound ABCA4 (EMD-XXXX, PDB XXX) | ATP-bound ABCA4 (EMD-XXXX, PDB XXX) |
|---|---|---|---|
| **Data collection and processing** | | | |
| Magnification | 130,000 | 130,000 | 130,000 |
| Voltage (kV) | 300 | 300 | 300 |
| Electron exposure ($e^{-}$/Å$^2$) | 50 | 50 | 50 |
| Defocus range (μm) | -2.0 to -1.0 | -2.0 to -1.0 | -2.0 to -1.0 |
| Pixel size (Å) | 1.08 | 1.08 | 1.08 |
| Symmetry imposed | C1 | C1 | C1 |
| Initial particle images (no.) | 2,554,957 | 2,227,280 | 2,169,513 |
| Final particle images (no.) | 205,597 | 184,628 | 173,278 |
| Map resolution (Å) | 3.3 Å | 3.4 Å | 3.3 Å |
| FSC threshold | 0.143 | 0.143 | 0.143 |
| Map resolution range (Å) | 3.0-5.0 Å | 3.0-5.0 Å | 3.0-5.0 Å |
| **Refinement** | | | |
| Initial model used (PDB code) | 5XJY | | |
| Model resolution (Å) | 3.3 Å | 3.4 Å | 3.3 Å |
| FSC threshold | 0.143 | 0.143 | 0.143 |
| Map sharpening *B* factor (Å$^2$) | -119 | -114 | -114 |
| Model composition | | | |
| Nonhydrogen atoms | 15,703 | 15,820 | 15,203 |
| Protein residues | 2,007 | 2,006 | 1,966 |
| Ligands | 19 | 21 | 21 |
| *B* factors (Å$^2$) | | | |
| Protein | 55.71 | 61.35 | 54.44 |
| Ligand | 63.93 | 73.81 | 82.16 |
| R.m.s. deviations | | | |
| Bond lengths (Å) | 0.008 | 0.009 | 0.010 |
| Bond angles (°) | 0.968 | 0.970 | 1.002 |
| **Validation** | | | |
| MolProbity score | 2.20 | 2.13 | 2.04 |
| Clashscore | 10.04 | 9.16 | 8.40 |
| Poor rotamers (%) | 0.00 | 0.00 | 0.00 |
| Ramachandran plot | | | |
| Favored (%) | 83.91 | 86.18 | 88.39 |
| Allowed (%) | 15.78 | 13.57 | 11.20 |
| Disallowed (%) | 0.30 | 0.25 | 0.41 |
